# Estimating seed dispersal distance: a comparison of methods using animal movement and plant genetic data on two primate-dispersed Neotropical plant species

**DOI:** 10.1101/528448

**Authors:** Tiziana A. Gelmi-Candusso, Ronald Bialozyt, Darja Slana, Ricardo Zárate Gómez, Eckhard W. Heymann, Katrin Heer

**Affiliations:** Verhaltensökologie & Soziobiologie, Deutsches Primatenzentrum – Leibniz-Institut für Primatenforschung, Kellnerweg 4, 37077 Göttingen, Germany; Naturschutzbiologie, Phillips-Universität Marburg, Karl-von-Frisch-Str. 8, 35043 Marburg, Germany; Nordwestdeutsche Forstliche Versuchsanstalt, Grätzelstr. 2, 37079 Göttingen; Instituto de Investigaciones de la Amazonía Peruana (IIAP), Av. José A. Quiñones km. 2.5, Iquitos, Perú

**Keywords:** parentage analysis, individual-based modelling, animal movement, zoochory, seed coats, seed dispersal curve, *Parkia panurensis*, *Leonia cymosa*, tamarins, animal behaviour

## Abstract

1. Seed dispersal distances (SDD) critically influence the survival of seedlings, spatial patterns of genetic diversity within plant populations and gene flow among plant populations. In tropical forests, a large percentage of seeds is dispersed by animals, and their foraging behaviour and movement patterns determine SDD. Direct observations of seed dispersal events by animals in natural plant populations are mostly constrained by the high mobility and low visibility of their vectors. Therefore, diverse alternative methods are used to estimate SDD, but direct comparisons of these approaches within the same seed dispersal system are mostly missing.
2. In this study, we take advantage of two plant species with different life history traits, *Leonia cymosa* and *Parkia panurensis* that are exclusively dispersed by two tamarin species, *Saguinus mystax* and *Leontocebus nigrifrons* (Callitrichidae) at our study site in the Peruvian Amazon. We compare SDD estimates obtained from direct observations, genetic identification of mother plants from seed coats, parentage analysis of seedlings, and modelling approaches, including the combination of movement data and gut passage times and individual-based modelling.
3. We detect differences between SDD estimates that can be linked to the processes relevant at different phases of the seed dispersal loop covered by the respective approaches. Despite these differences, SDD estimates for *P. panurensis* are consistently lower than for *L. cymosa* which is likely related to differences in fruit characteristics and fruit abundance, factors that influence gut passage time, foraging behaviour and movement patterns of the tamarins.
4. Our comparisons allow setting SDD estimates from studies using different methodological approaches into the seed dispersal loop context, thus improving comparability of methodologically different studies and method applicability.

## Introduction

Seed dispersal provides the spatial template for subsequent processes that ultimately result in the recruitment of new individuals in plant populations. It impacts seed survival (Janzen, 1970; Connell, 1971; Schupp, Jordano, & Gómez, 2010; Schupp & Jordano, 2011; Steinitz, Troupin, Vendramin, & Nathan, 2011), determines gene flow within and among populations (Cain, Milligan, & Strand, 2000; Ran Nathan et al., 2008; He, Lamont, Krauss, & Enright, 2010), fosters habitat connectivity (Culot, Lazo, Huynen, Poncin, & Heymann, 2010; Lindsell, Lee, Powell, & Gemita, 2015; Ripperger, Kalko, Rodriguez-Herrera, Mayer, & Tschapka, 2015) and enhances the probability of survival of populations under anthropogenic pressure (Snyder, 2011; McConkey et al., 2012; Ruxton & Schaefer, 2012; Abedi-Lartey, Dechmann, Wikelski, Scharf, & Fahr, 2016). Measuring the seed dispersal distance (SDD), i.e. the distance between the source plant of a seed and its deposition site, is crucial for determining the spatial dimension of the seed shadow and predicting the outcomes of the processes following seed dispersal (Nathan & Muller-landau, 2000; Nathan, Klein, Robledo-Arnuncio, & Revilla, 2012).

Different approaches have been employed to estimate SDD and seed shadows, each having its specific benefits and limitations (Table 1). Naturally, the method of choice depends on the specific study system and the resources available. Nathan et al. (2012) describe different methods to estimate seed dispersal kernels, comparing limitations and uses, however, direct comparisons between methods within the same seed dispersal system are scarce. Mise *et al.* (2016) compared seed dispersal distances estimates for racoon dogs (*Nyctereutes procyonoides*) using the bait-marker method against the combination of movement data and gut passage, and found comparable results when data was collected from the same region. For the Neotropical legume *Parkia panurensis*, the matching of genotypes of seed coats to potential maternal trees was confirmed with SDD obtained through direct observation of seed dispersal vectors (Heymann et al., 2012). Spatially explicit modelling based on movement data from the same vectors, but from an independent study, further supported these findings (Bialozyt, Flinkerbusch, Niggemann, & Heymann, 2014). Using different methods for determining SDD within the same study system allows for an evaluation of the comparability between results from different methods, most important when deriving conclusions for seed dispersal systems based on different studies applying different methods. Using primate-dispersed plants for such a comparison is particularly useful since primates, once habituated, can be followed for direct observations of seed dispersal events. Thereby, the position of feeding plants and seed dispersal sites can be recorded, an approach rarely feasible with birds, bats, and other highly mobile vectors. Finally, this comparison provides valuable information on which method to choose when direct observations are not possible because the target plant species of a study system fails to produce fruits. Such phenological changes may become increasingly more frequent as a consequence of global climate change (Cleland, Chuine, Menzel, Mooney, & Schwartz, 2007; Abernethy, Bush, Forget, Mendoza, & Morellato, 2018) and make it increasingly harder to plan field studies on seed dispersal. For example, in our specific case, after years of high fruiting yield, the population of *L. cymosa* at our study site produced a deficient number of fruits during our sampling years, leading us to rely on alternative methods for estimating SDD.

**Table 1.** Comparison of the advantages and disadvantages of different approaches for estimating seed dispersal distance.

| Method | Data/material required | Advantages | Limitations | References |
| --- | --- | --- | --- | --- |
| Observed seed dispersal events (OSD) | Location of seed sources (food plants) and seed deposition sites | Simple, direct SDD estimate from locational data<br>Sampling of seed source data can be restricted to those plants that were visited by the vector | Seed source can only be reliably identified if vectors do not feed on other individuals of the same plant species between first feeding on that species and seed deposition<br>Seed dispersal vectors may be complicated to follow | Yumoto, Kimura, & Nishimura, 1999; Stevenson, 2000; E. V. Wehncke, Hubbell, Foster, Dalling, & Hubbell, 2003; Tsuji, Yangozene, & Sakamaki, 2010; Valenta & Fedigan, 2010; Heymann et al., 2017 |
| Genotyping of seed coats (GSC) | Seeds with seed coats<br>Location of all potential source plants and seed deposition sites<br>Tissue samples from adult plants<br>Nuclear markers (e.g. SSRs) | Accurate and reliable identification of source plants<br>Can be used for accumulation sites and seed traps.<br>No need to follow vectors | High sampling effort for seeds and adults needed<br>Finding seeds may become impractical if vectors have large ranging areas<br>If multiple vectors disperse seeds of a plant species, the contribution of each vector to SDD cannot be estimated<sup>1</sup> | Dow & Ashley, 1996; Godoy & Jordano, 2001; Grivet, Smouse, & Sork, 2005; Smouse, Sork, Scofield, & Grivet, 2012 |
| Parentage analysis of seedlings (PAS) | Location of potential source plants and seedlings<br>Tissue samples from adult plants and seedlings<br>Nuclear and/or chloroplast markers (e.g. SSRs) | Measures the outcome of seed dispersal<br>No need to follow vectors | Measures the outcome of seed dispersal and includes undispersed individuals<br>High sampling effort of adults needed<br>Accurate identification of source plants with nuclear markers only in dioecious plants | Hamrick & Trapnell, 2011; Choo, Juenger, & Simpson, 2012 |
| Modelling of SDD based on daily travel path and gut passage time (CMG) | Vector movement data (observation, radio-tracking, GPS-tracking)<br>Gut passage time estimate | Estimates potential SDD for single vectors, even within complex seed dispersal networks<br>Does not require finding dispersed seeds and sampling source trees | Includes movements beyond post-feeding events<br>Unless vectors can be observed from feeding to seed deposition in nature, estimating gut passage times requires capturing and experimental feeding | Murray, 1988; Holbrook & Smith, 2000; Westcott et al., 2005; Fuzessy, Janson, & Silveira, 2017 |
| Individual-based modelling of SDD (IBM) | Focus plant fruiting characteristics<br>Focus plant locations<br>Additional fruiting tree locations<br>Vector movement data (observation, radio-tracking, GPS-tracking) | Estimates potential seed dispersal for single vectors, even within complex seed dispersal networks<br>Does not require finding dispersed seeds | Unless vectors can be observed from feeding to seed deposition in nature, estimating gut passage times requires capturing and experimental feeding | Pakeman, 2001; Morales & Carlo, 2006; Russo, Portnoy, & Augspurger, 2006; Couvreur et al., 2008; Levey, Tewksbury, & Bolker, 2008; Uriarte et al., 2011; Bialozyt, Flinkerbusch, et al., 2014 |
| Seed-tracking (not used in this study) | Location of source plants or of experimental seed source and seed deposition | Controlled experiment | Experimental approach may not fully capture the variation of the seed dispersal process caused, e.g. by variable patterns of vector movements | Levey & Sargent, 2000; Chauvet, Feer, & Forget, 2004; Pons, Pausas, & Pausas, 2007; Hirsch, Kays, & Jansen, 2012; Jansen et al., 2012; Sork, 2016 |

In this paper, we compared different approaches for estimating SDD and seed dispersal curves. We used data from the previous studies on *P. panurensis* (Heymann et al., 2012; Bialozyt, Flinkerbusch, et al., 2014) where SDD had already been estimated based on direct observation of seed dispersal events, maternal identification through genotyping of seed coats and individual-based modelling, and added SDD estimates based on two additional approaches. We replicated these five approaches for *L. cymosa*, which is dispersed by the same vectors as *P. panurensis*, but for which we obtained only a few direct observations due to unexpected low fruit production. Therefore, for both species, we compare SDD estimates based on five methods: (1) <u>o</u>bserved <u>s</u>eed <u>d</u>ispersal events (OSD), (2) seed dispersal estimates from maternal identification through genotyping of <u>s</u>eed <u>c</u>oats (GSC), (3) <u>p</u>arentage <u>a</u>nalysis of <u>s</u>eedlings (PAS), (4) modelling of SDD through a <u>c</u>ombination of <u>m</u>ovement data with gut passage times (CMG), and (5) simulation of seed dispersal by <u>i</u>ndividual-<u>b</u>ased <u>modelling</u> (IBM). Apart from the methodological comparison, our analysis also allows comparing the effect of plant traits on SDD without the potentially confounding effect of different dispersal vectors.

## Materials and Methods

### Study site

Data for this paper were collected at the Estación Biológica Quebrada Blanco (EBQB), located at 4°21’S 73°09’W in north-eastern Peruvian Amazonia. For details of the study area, see Heymann (1995) and Culot et al. (2010).

### Plant species

We use data from two different plant species, *Leonia cymosa* (Violaceae) and *Parkia panurensis* (Fabaceae) for our comparisons. *Leonia cymosa* is an understory tree and occurs at an adult population density of 5 and 11 individuals/ha within the home ranges of the investigated tamarin groups at EBQB. It produces berries with 2-7 seeds embedded in a fibrous pulp. During weekly counts, 25-125 fruits were present per tree in a previous study (Reinehr, 2010). Fruits ripen asynchronously within and between trees throughout several weeks, up to three months. Feeding visits by tamarins vary between <1 and 10 min (mean: 1.9 min; SE: 0.1), with generally only 1-2 (maximum 5) tamarins feeding in a single tree (Reinehr, 2010). On average tamarins visit 7.5 trees per day (range: 1-14 and generally feed in several *L. cymosa* individuals in a row (Reinehr, 2010). This “trap-lining” (Garber, 1988) makes it challenging to assign seeds to a source tree during the observation of seed dispersal events.

*Parkia panurensis* is a canopy tree and occurs at a density of around one adult/ha. It produces pods with 15-25 seeds surrounded by an edible sticky gum (Hopkins, 1986) Within a single tree, ripe pods may be present for up to 12 weeks, within the population for up to ca. 4 months. Feeding visits of tamarins’ range between 3 and 46 min (mean: 11.8 min; SE: 0.7), depending on the number of ripe pods available at a given moment, and generally, all group members are feeding simultaneously in a single tree (Heymann, pers. obs.; Feldmann, 2000). Tamarins may visit the same tree up to four times per day, and there is no trap-lining. Therefore, seeds can very often be reliably assigned to a single source tree (Heymann et al., 2012).

#### Seed dispersal vectors

At EBQB, the seeds of both plant species are exclusively dispersed by two species of tamarin monkeys, *Saguinus mystax* and *Leontocebus nigrifrons* (Callitrichidae). These primates live in groups of 3-12 individuals and form mixed-species troops in which members from both species move through a shared home range in a highly coordinated way (Heymann & Buchanan-Smith, 2007). Home range size varies between ca. 25 and 50 ha and mean daily path length (i.e. the length of the route travelled from sleeping site to sleeping site) ranges from 600 - 3,000 m (mean: 1,700 m; Smith, 1997). Daily travel paths are not linear but have variable shapes (e.g., concentric, meandering; see Supplementary Information 1 for examples).

#### Collection of observational data

Tamarin groups at EBQB are well-habituated to the presence of human observers and can be observed at close range. Data on feeding and seed dispersal of *L. cymosa* and *P. panurensis* are available from studies by Knogge (1999), Culot (2009) and Heymann et al. (2012). They include location and time of feeding and defecation of both tamarin species. Defecations containing one or more seeds were defined as seed dispersal events (Knogge & Heymann, 2003). Movement data of tamarin groups, independent of seed dispersal events, were recorded for a total of 62 days, with a mean of 7.7 ± 2.8 days per month from December 2012 to July 2013. Position data were recorded with GPS devices or previous to GPS devices (1999) determined in reference to the 100 m x 100 m grid system and in reference to previously marked and mapped trees at EBQB.

#### Sampling of plant material

We collected leaf samples from *L. cymosa* seedlings (height <100 cm), juveniles (100-250 cm) and adult plants (>250 cm) in 2014. We exhaustively sampled all plants in a total of 25 quadrats of 50 m × 50 m) within the home range of tamarin group 1 (12 quadrats ~ 15 % of home-range area) and 2 (13 quadrats ~ 15 % of home-range area). Quadrats were located at the crossings of the trail system that spans the study site (Figure 1). To increase the number of adults, we additionally sampled along 53 transects of 15 m x 100 m in Group 1 and additional quadrats on remaining path intersections in Group 2. Finally, when the trees were fruiting in 2016, we collected seeds of *L. cymosa* that were excreted by tamarins, stored them in saline solution and recorded the location of deposition. For the genetic analysis, we either dried the sampled leaf tissue on silica gel beads or soaked Whatman FTA Plantsaver cards (GE Healthcare Lifesciences, Chicago, IL, USA) with smashed leaf material (see Supporting information S1).

**Figure 1.**
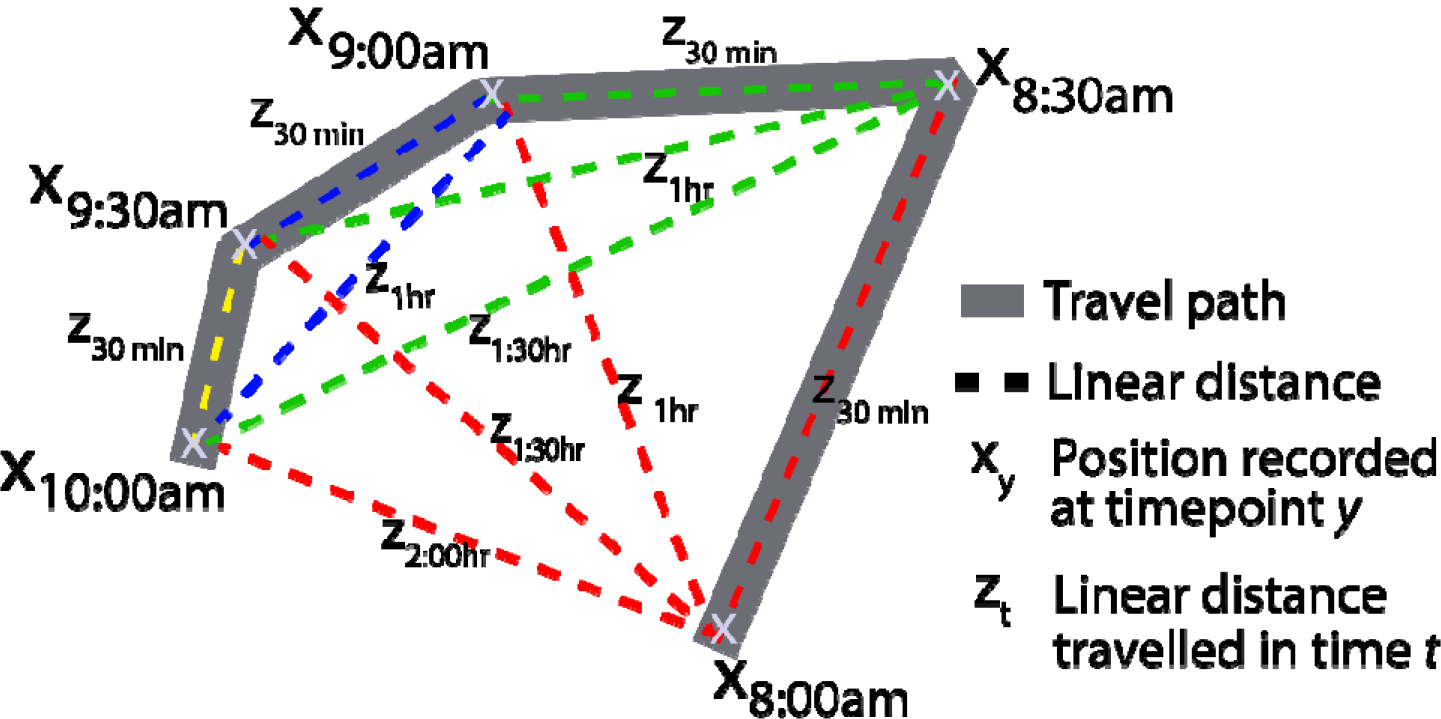
Illustration of the CMG method. Graphical example for the calculation of daily travel path for the modelling of SDD through a <u>c</u>ombination of <u>m</u>ovement data with gut passage times (CMG). To obtain a series of linear distances (dashed lines), we calculated the linear distances between scan points (X) that were recorded every 30 min throughout the day along the travel path of the tamarins. This way, we obtained a series of distances for different time intervals from 30 min up to 9 h for each day.

A full inventory of adult *P. panurensis* (dbh >20 cm) was done within the home range of tamarin Group 1 (Bialozyt, Luettmann, et al., 2014). Geographic coordinates were recorded with a GPS device (Garmin GPSMap 76CSx). Seedlings (height <1.3 m) and juveniles (height >1.3 m and dbh <20 cm) of *P. panurensis* were sampled by randomly overlaying a 50 m × 50 m grid over a map of the home range of tamarin Group 1. Intersections of this grid were taken as central points for 50 m × 50 m quadrants were seedlings and juveniles were sampled exhaustively.

#### Genotyping with microsatellite markers

To prepare the leaf samples of seedlings and adult trees for DNA extraction, we ground the leaves using a Retsch mill (Haan, Germany). Seeds were first rehydrated at room temperature, and then seed coats were carefully separated from the embryo. We dried the seed coats (i.e. maternal tissue) on filter paper and ground them in the Retsch mill. DNA was extracted from ground leaves and seed coats following the modified CTAB protocol with ATMAB (Dumolin, Demesure, & Petit, 1995). DNA of each sample was genotyped using eleven nuclear microsatellites for *L. cymosa* following the protocol described in Supplementary data 2. For *P. panurensis* we used genotype information of 33 adults and 85 seedlings and 92 seeds that were previously genotyped with nine nuclear microsatellite markers (Heymann et al., 2012).

#### Data analysis for estimates of seed dispersal distances

##### I. SDD from observed seed dispersal events (OSD)

For each observed dispersal event, we determined the SDD as the linear distance between the location of feeding and defecation.

##### II. Maternal identification from genotyping of seed coats (GSC)

The seed coat is of maternal origin, and thus a precise match between the genotype of a seed coat and potential mother trees would identify the right mother. Genotypes from seed coats were matched to adult genotypes using GenAlex v. 6.501 with no mismatches allowed (Peakall & Smouse, 2006). Euclidean distance between the source tree and the dispersed seed was determined based on the recorded UTM coordinates.

##### III. Parentage analysis of seedlings (PAS)

We used the CERVUS software v3.0.7 (Slate, Marshall, & Pemberton, 2000) to determine the potential parents for the genotyped seedlings in our study area. We selected strict confidence intervals (95%) for the parental analyses using maximum likelihood framework. We run the preliminary simulation with the following parameters settings: number of candidate parents was set to 202 for *L. cymosa* and 33 for *P. panurensis*, proportion sampled set to 0.15 for *L. cymosa* based on sampling areas in relation to the total home range area and 0.99 for *P. panurensis* were we are confident that all adult trees in the tamarins′ home range were sampled. Finally, the genotyping error was determined as 0.01 using GenAlex, and we only included individuals with a minimum of six typed loci for *L. cymosa* and five for *P. panurensis*. Afterwards, we determined the Euclidean distance between the resulting parents and offspring.

Given that maternal sources of seedlings and juveniles were unknown in *L. cymosa* and *P. panurensis*, we assumed that both parents could be either mother or father to avoid potential bias. Following this assumption, we used all possible parent-seedling combinations to calculate a mean SDD and density distance kernels. As observations of seed dispersal events showed that seed dispersal distances by tamarins do not exceed 709 m (N=1884; Knogge, 1998) which corresponds to the diameter of a tamarin home range, we excluded seedling-parent pairs at distances > 709 m from this analysis assuming that this is instead a pollen `source` than a seed source.

##### IV. Combination of movement data with gut passage time estimate (CMG)

To determine seed dispersal distances based on daily travel paths of dispersal vectors we modified the approach used by Murray (1988). Linear distances between scan points were calculated for each daily travel path, considering the time interval between each pair of scan points (**Figure 1**). We created a function in R, *linear.distances*(), (see supporting information S2) to automatically extract the information from movement data collected in 2013. From the movement data, we obtained a series of linear distances for time intervals from 30 min up to 9 hours for each day. In contrast to Murray′s approach, we did not limit the analysis to scan points after visits to food plants. Thus, our method is also applicable under conditions where no information on the timing of feeding is available, e.g. in cases where animals are tracked remotely.

In a next step, we only considered linear distances within the time interval of gut passage for the seeds of each species. We estimated gut passage time as the time lag between feeding and defecation (from data collected by Knogge, 1998 and Culot, 2009). Since resting times can increase gut passage time without increasing seed dispersal distance, we used a conservative range with the upper limit within 80% confidence interval (CI) and the lower limit within 5% CI of passage times, calculated with the function *npquantile*() from the “np” package (Hayfield and Racine, 2008) for skewed distributions. The resulting gut passage time estimate for *Parkia panurensis* was 0:30-4:00 hrs (N=196)*. For L. cymosa* we had only a few observation data and used the minimum and maximum value observed, resulting in a gut passage time estimate of 2:00-4:00 hrs (N=3).

To account for monthly variation of travel path length, we only considered the fruiting periods for each species (May - July for *P. panurensis* and Mar to May for *L. cymosa*). With this subset of potential seed dispersal events, we plotted the density distance kernel using the *bkde*() function from the “KernSmooth” package following suggestions by Deng & Wickham (2011). For each method, we decided the bandwidth for the density curve based on the function *density*().

##### V. Individual-based modelling of seed dispersal events (IBM)

Individual-based modelling (IBM) provides another approach to assess SDD. In such a model, the tamarins move around and eat fruits in order to maintain homeostasis and as a result disperse seeds after a predefined gut passage time somewhere within their home range. Bialozyt et al. (2014a) developed such a model *P. panurensis* at the same study site and for the same tamarin home range. In their model, they used three different tree types: feeding, resting and sleeping trees. In the model, only those *P. panurensis* trees in the home-range area that were used as actual feeding trees during the observation period in 2008 were considered and – for the purpose of the simulation – it was assumed that there were no other species used for feeding. This assumption was valid for these simulations because the data collection had been carried out during a timespan when *P. panurensis* was nearly the exclusive fruit source for the tamarins.

In the current paper, we adjusted the previous model for the seed dispersal scenario of *L. cymosa*. Thus, we needed to adjust four critical aspects. First, since *L. cymosa* is never the only fruit source available in this area, we needed to add other species as fruit source to allow for enough energy input during the daily routine of the tamarins. We used the other species of feeding trees observed during *L. cymosa*’s fruiting season in 2013 as additional fruit sources. Furthermore, not all *L. cymosa* trees fruit yearly; therefore, we used the subset of *L. cymosa trees* (N = 8) observed that same year. Second, *L. cymosa* contains 415-642 mg of soluble sugars per g dry matter, whereas *P. panurensis* contains 811 mg/g (Pfrommer, 2009; Peres, 2010); therefore, we adjusted the mean energy level provided by the trees in the simulation model. Third, different time intervals in feeding trees for a single feeding event were implemented for *P. panurensis* and *L. cymosa* (Table S4.5) to reflect the respective fruit crop sizes and the resulting shorter feeding times in *L. cymosa*. Fourth, we adjusted gut passage time for *L. cymosa* according to field observations of seed dispersal events reported by Knogge (1999) and Culot (2009). All other parameters were kept at values of the *P. panurensis* simulation (Bialozyt et al. 2014a).

Simulations of daily movements were carried out for 200 days to get enough *L. cymosa* seed dispersal events. We then determined the Euclidian distance between dispersed seeds and their mother trees. Details on the simulation model are presented in Bialozyt et al. (2014a), and changes to the original model are provided in supporting information S4 (Table S4.1).

#### Statistical analysis of seed dispersal distance

We estimated mean SDD values for each method through bootstrapping distance values (n = 10,000 resamplings), using the *boot_mean*() function from the “boot” package in R (Davison & Hinkley, 1997; Canty & Ripley, 2017). We evaluated differences between methods and species with the non-parametric Kruskal-Wallis test using the *kruskal.test*() function from the stats package in R (R Core Team, 2018). We did further posthoc comparisons with the non-parametric multiple comparison test and Bonferroni corrections, using the *pairwise.wilcox.test*() function, from the stats package in R (R Core Team, 2018).

To estimate seed dispersal curves, we determined the empirical frequency distribution (i.e. density distance kernels) of dispersal distances for each method by adjusting a non-parametric function (smooth spline curve) and its confidence envelope estimated by bootstrapping (n = 100 resamplings) using the *mykernel*() function (Jordano, 2016). Bandwidth size was calculated with the function *density*() from the “stats” package (R Core Team, 2018).

Finally, to compare seed dispersal curves between methods we estimated the probability distribution of all methods using the *stat_ecdf*() function from the “ggplot2” package in R (Wickham, 2016). Subsequently, we tested differences between the empirical cumulative distribution functions of each method with the two-sample Kolmogorov-Smirnov test, which is sensitive to differences in both location and shape of the cumulative distribution function. For the Kolmogorov-Smirnov test, we used the *ks.test*() function from the package “stats” in R (R Core Team, 2018).

## Results

### Comparison among methods

Depending on the method used, mean SDD estimates range between 158 and 201 m for *P. panurensis* and between 178 and 318 m (30% difference) for *L. cymosa*. Overall, methods varied significantly in the resulting SDD estimates (*P. panurensis*: H_(4)_ = 13.7, p = 0.009; *L. cymosa*: H_(4)_ = 17.3, p = 0.002). Specifically, Wilcoxon pairwise comparisons revealed that SDD estimates from IBM were significantly higher than those from OSD and GSC, and SDD estimates from CMG were significantly higher than those from GSC, in *P. panurensis* (Fig. 2a) and SDD estimates from PAS were significantly lower than those from GSC, CMG and IBM in *L. cymosa* (Fig. 2b).

**Figure 2.**
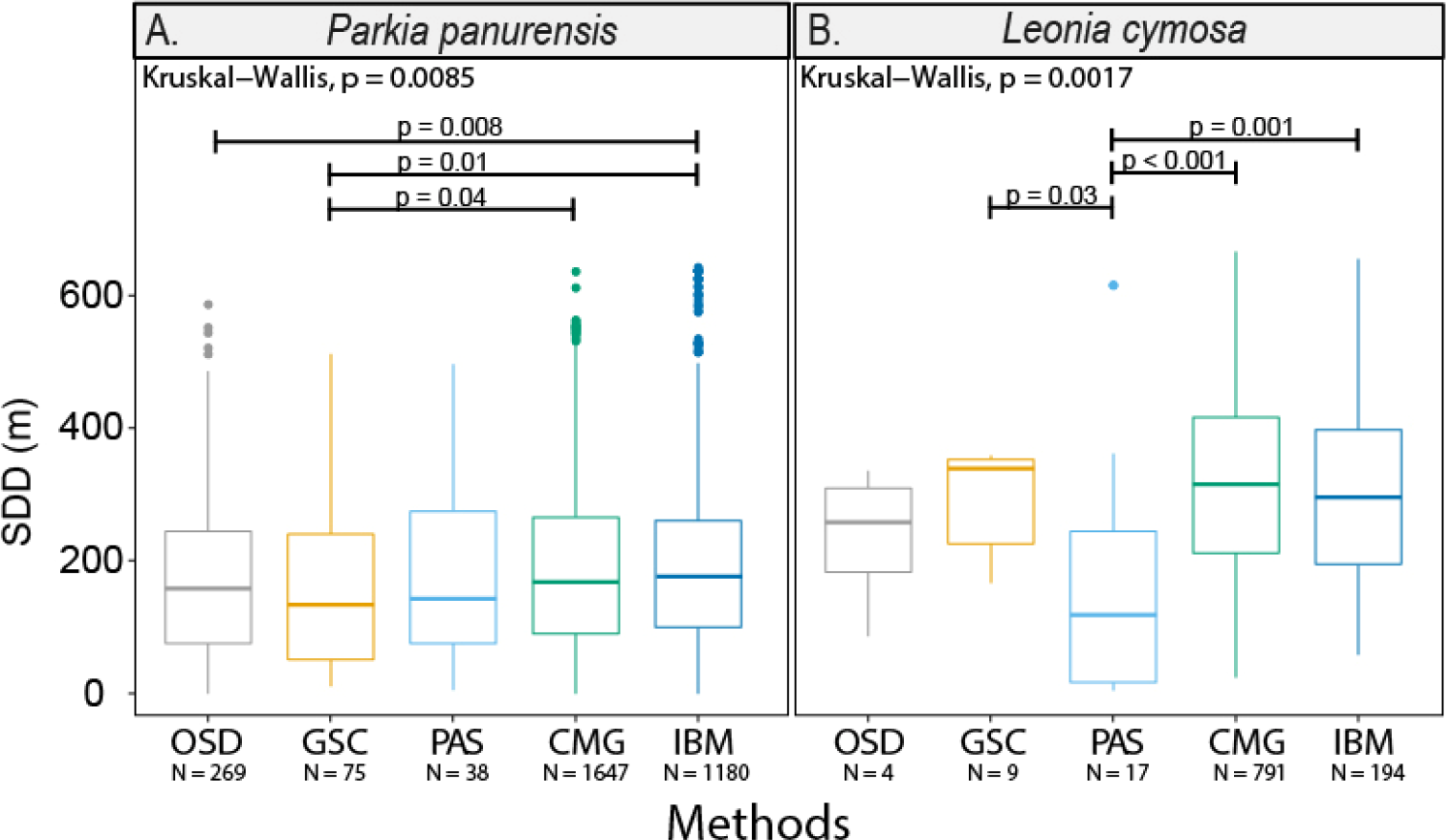
SDD estimates based on the five methods used in this study: observed seed dispersal events (OSD), genotyped seed coats (GSC), parental analysis of seedlings (PAS), combination of movement data and gut passage (CMG), and individual-based modelling (IBM) for (a) *Parkia panurensis* and (b) *Leonia cymosa*. Horizontal lines represent medians, boxes the 25-75% quartiles, dots are outliers. Bars above the boxplots indicate differences among methods based on a Kruskal Wallis test and multiple pairwise comparisons with Wilcoxon rank sum test.

For *P. panurensis*, all methods except for IBM provide a strongly significant right-skewed distribution of SDD OSD, GSC, and CMG (Kolmogorov-Smirnoff test: p=0.02, p=0.03, p=0.002, respectively) (Fig. 3a). Furthermore, in *P. panurensis*, the cumulative SDD curves of all methods converge at low distances, (Fig. S3.4a). In *L. cymosa*, the shape of the SDD distributions was highly variable on results from methods executed with small sample number, and only the curve based on PAS approaches a right-skewed pattern while the others show a trend towards normality. PAS curve was significantly more left-skewed than GSC, CMG, and IBM (Kolmogorov-Smirnoff test, p=0.03, p<0.001, and p<0.001, respectively) (Fig.3b)

**Figure 3.**
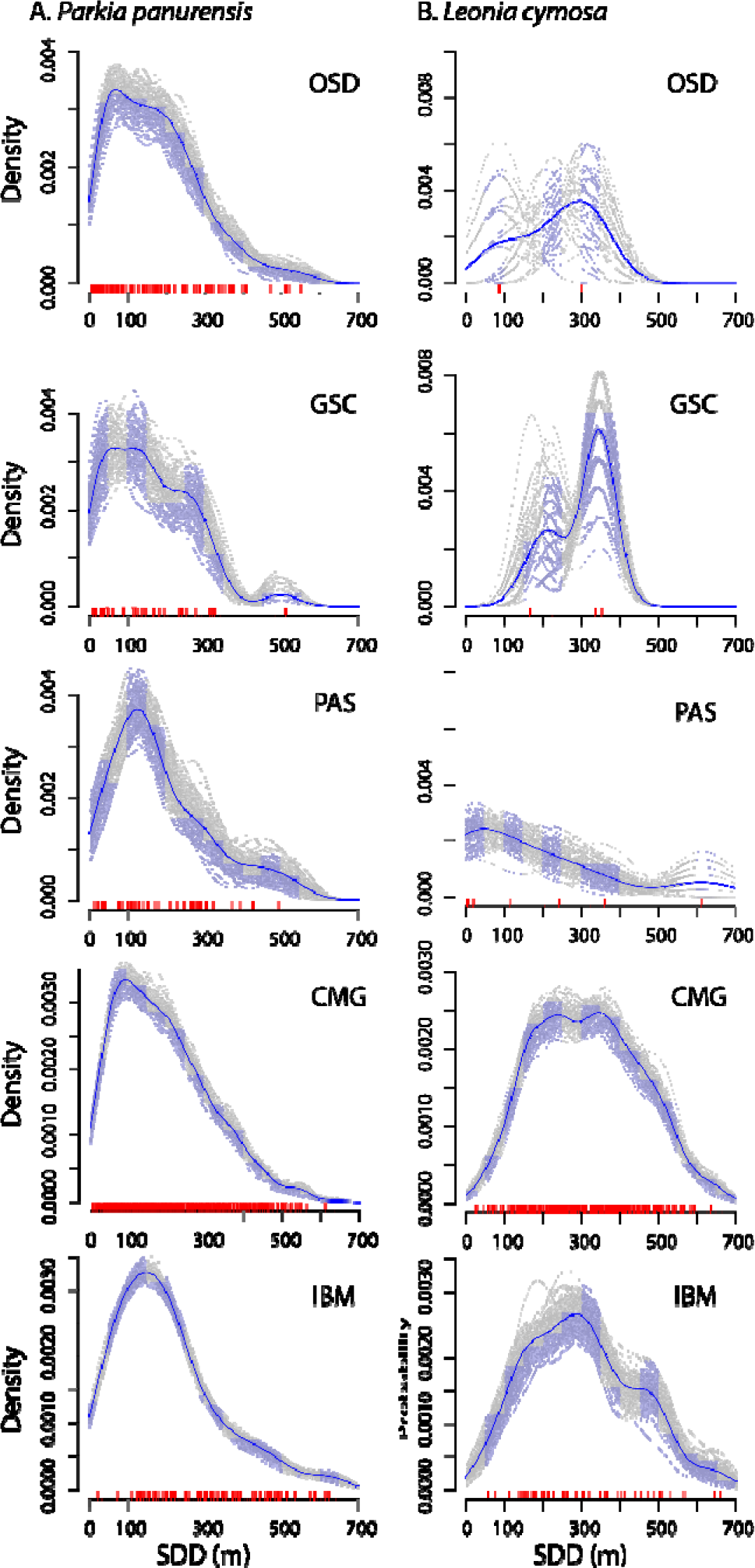
Distribution of seed dispersal distances for the five methods used for *Parkia panurensis* (A) and *Leonia cymosa* (B). The figures show for each method, the density of dispersal events within the distance class (blue bars), a nonparametric smoothing spline fit to the empirical distance distributions (blue lines) together with bootstrapped estimates (grey lines). Red vertical bars along the x◻Jaxis represent each observed dispersal event.

### Comparison between species

SDD estimates varied significantly between *P. panurensis* and *L. cymosa* (H_(4)_ = 557.5, p < 0.001). Specifically, Wilcoxon pairwise comparisons revealed that estimates obtained through GSC, CMG and IBM differed significantly with higher SDD for *L. cymosa* (Fig S3.5).

## Discussion

Our analyses revealed that different methods resulted in different SDD estimates for two Neotropical rainforest tree species. Each method refers to different processes within the “seed dispersal loop” described by Wang and Smith (2002, Fig 4), which could explain part of the differences. For example, while OSD estimates SDD after step 2 of the seed dispersal loop including the resulting seed deposition, CMG measures SDD at step 2 of the seed dispersal loop without including seed deposition, therefore giving only an estimate of all potential seed dispersal distances. IBM integrates over steps 1 and 2, providing a range of simulated seed dispersal distances based on possible case scenarios deriving from only these two steps. PAS, instead, estimates SDD after step 4 (and eventually step 5). Therefore, it includes also germination success and post-dispersal processes, which might lead to lower or higher estimates according to plant-specific density-dependent mortality (Fig. 4).

**Figure 4.**
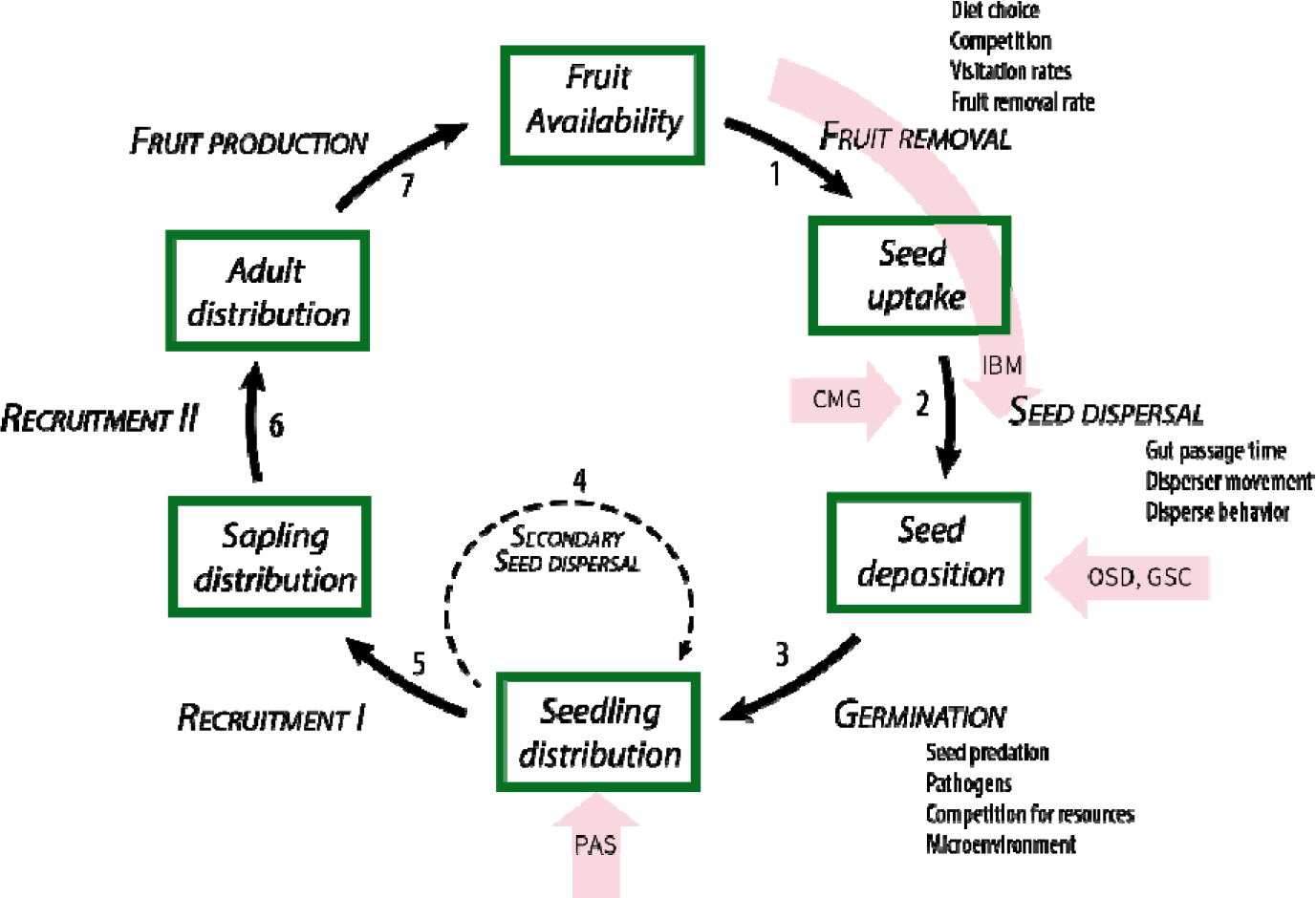
Seed dispersal loop as (modified from Wang and Smith 2002) showing which processes or steps of the dispersal loop are integrated by each method we used for estimating SDD

However, different positions within the seed dispersal loop may not explain the fact that different methods provide lower vs higher estimates in an inconsistent way for different species. Here, we explore how these inter-specific inconsistent differences between methods could be related to different life histories, species-specific interactions with seed dispersers and post-dispersal processes in the two plant species. In the case of *P. panurensis*, differences between methods only concerned the median SDD, but not the overall distribution of SDD which was consistent and showed a right-skewed pattern that seems to be typical for SDD distributions (Clark, Silman, Kern, Macklin, & HilleRisLambers, 1999; Nathan et al., 2008). In contrast, for *L. cymosa* not only median SDD differed between methods, but also the overall SDD distributions, where most curves deviated from the right-skewed pattern showing normality and even slight bi-modality, except for the curve based on PAS. Even though for the curves based on OSD and GSC, the minimal sample size could play a role on this difference, a right-skewed curve for the only method including germination success could potentially imply a high germination success of undispersed seeds near source trees.

Our results indicate that comparisons between studies need to carefully evaluate the differences among methods for estimating SDD. On the other hand, a combination of methods can be used to obtain more information on the seed dispersal system, when considering the different steps of the seed dispersal loop taken into account by each method.

Reliable estimates of gut passage time are crucial for modelling SDD with the CMG method. In both tamarin species, gut passage times show considerable variation within and between plant species (Knogge, 1999). Therefore, the higher the sample size of gut passage times, the more reliable the results obtained from this method. However, our results show that even a small sample size for gut passage times results in similar SDD estimated obtained from CMG and IBM. An alternative to field observations of gut passage times are estimates derived from captive animals. A number of studies estimated gut passage times (or regurgitation times) from captive feeding experiments with one or few plant species and then extrapolated to other plant species to estimate SDD (Holbrook & Smith, 2000; Westcott, Bentrupperbäumer, Bradford, & McKeown, 2005; Holbrook, 2011; Abedi-Lartey et al., 2016). It remains to be determined how representative these results from captive animals are, as restriction of movements affects gut motility (Holdstock, Misiewicz, Smith, & Rowlands, 1970; Oettlé, 1991). For example, gut passage time of seeds increased by up to 80% with physical activity in mallards (Kleyheeg, van Leeuwen, Morison, Nolet, & Soons, 2015). Studies of primate seed dispersal are “privileged” as gut passage times can be determined from unconstrained, naturally moving animals (Bravo, 2009.; Knogge, 1999; Wehncke, Hubbell, Foster, & Dalling, 2003; Valenta & Fedigan, 2010) which is mostly not feasible in birds, and impossible for nocturnal bats. Yet, in these taxa, remote tracking of animals via GPS telemetry is becoming increasingly feasible with new and smaller tracking devices (Abedi-Lartey et al., 2016; McMahon et al., 2017; Oleksy, Giuggioli, McKetterick, Racey, & Jones, 2017) that allows tracking of small to medium-sized seed dispersers. These devices allow obtaining data of daily travel paths that will provide fine-tuned estimates of SDD using the CDG method if reliable data of gut passage time is available.

Despite the uncertainty of gut passage time in *L. cymosa* it is plausible to assume that gut passage times are longer for *L. cymosa* compared to *P. panurensis*. In the former, the fibrous pulp is firmly attached to the seeds, in contrast to the gelatinous exudate in the latter, and Knogge (1999) showed that seeds with fibrous pulp produce longer gut passage times than seeds with gelatinous pulp. So even if tamarin movement patterns were identical when feeding on these two species, the longer gut passage time would increase SDD. Also, movement patterns and foraging behaviour of tamarins differ when feeding in *L. cymosa* and *P. panurensis*. In the former, only a few individuals feed simultaneously in the same tree, and the same tree is rarely revisited on a given day (Reinehr, 2010). In the latter, the entire group feeds simultaneously, and the same tree can be repeatedly visited on the same day, which creates overlaps in the travel path. This overlap is reflected in the estimates based on IBM, which result in values comparable to the other methods for *P. panurensis* but not for *L. cymosa*.

In conclusion, our comparative approach allows for better comparability of SDD estimates from studies that employ only one or two of the five approaches in this study. We identified the stages of the seed dispersal loop considered by each method and highlight their advantages and constraints. Despite differences in SDD estimates of the compared methods, estimates were consistently lower for *P. panurensis* than for *L. cymosa.* We linked this to plant species-specific traits that influence gut passage time, foraging behaviour and post-feeding movement `patterns` of the tamarins.

## Acknowledgements

The authors thank C. Mengel, C. Glaschke and C. Schwarz for their support with laboratory work, and L.A. Garcia Ayachi, C. Flores Amasifuén and N. Shahuano Tello for assistance with sample and data collection in the field. Sampling and genotyping of *Leonia* were funded by the German Research Council (Deutsche Forschungsgemeinschaft, grants HE 1807/20-1, and HE 7345/1-1). The study was conducted under authorisation no. 0160-2014-MINAGRI-DFGGS/DGEFFS from the Peruvian Ministry of Agriculture.

## Data accessibility

All datasets and the codes for the CMG and IBM method are available as supplementary files and will be archived in *Dryad*.

## Author contribution

TG, EWH, and KH conceived and designed the analysis. TG and RZ, collected plant material and DS collected animal movement data. TG performed the genetic analysis, the *OSD*, *GSC*, *PAS, and CMG* analysis, the statistical analysis and prepared the figures. RB performed the *IBM analysis*. TG, EWH, and KH wrote the manuscript and TG, DS, RZ, RB, EWH, and KH revised the final version of the manuscript.

