## Supplementary Information for "Estimating seed dispersal distance: a comparison of methods using animal movement and plant genetic data on two primate-dispersed Neotropical plant species"

**Supporting Information – S1**

**Characterization of 11 microsatellite (SSRs) loci for *Leonia cymosa* and their compatibility with 2 congeneric species: *L. glycicarpa* and *L. crassa*.**

Microsatellite development and characterization

Leaf samples of *Leonia cymosa* were collected at Estación Biológica Quebrada Blanco (EBQB) and in the periphery of Allpahuayo-Mishana (AM) in north-eastern Peruvian Amazonia. Samples of *L. glycicarpa* and *L. crassa* were only collected at AM. Location and description of vouchers are described in Table 1.

Table 1 Description of specimen locations from sampled individuals

| Species | Voucher specimens | Collection locality | Geographic coordinates |
| --- | --- | --- | --- |
| *L. cymosa* | TA Gelmi-Candusso 025022 EBQB (AMAZ) | Estacion Biologica Quebrada Blanco (EBQB), Iquitos, Peru | 4° 21’ S,  73° 09’ W |
| *L. cymosa* | R Zarate 20272 AM (HH) | Alpahuayo-Mishana (AM), Iquitos, Peru | 4° 29' S,  73° 35' W |
| *L. crassa* | R Zarate 20273 AM (HH) | Alpahuayo-Mishana (AM), Iquitos, Peru | 4° 29' S,  73° 35' W |
| *L. glycicarpa* | R Zarate 20271 AM (HH) | Alpahuayo-Mishana (AM), Iquitos, Peru | 4° 29' S,  73° 35' W |

We stored leaves collected by drying them in silica gel beads (Sigma-Aldrich, St. Louis, MO, USA) or pressed into FTA Plantsaver cards (GE Healthcare, Chicago, IL, USA) and stored seeds collected on saline solution. We extracted DNA from dried leaf tissue and seed coats by homogenizing 100mg of each sample using a Retsch shaking mill (Retsch, Hilden, Germany) and subsequently following the ATMAB-based protocol (Dumolin et al., 1995) with an additional final treatment with 0.5 µl RNase at 37°C for 30 min. For leaf samples stored in FTA plantsaver cards, we extracted DNA following the company’s protocol and adding a final 5 min incubation step in TE buffer (TRIS, EDTA) at 95°C for obtaining solute DNA. DNA concentrations were measured using the NanoDrop 3300 (Thermo Fisher Scientific, Waltham, MA, USA). Both methods of storage for leaves yielded identical genotyping results when used on the same samples (unpub. data).

Microsatellites markers were developed by Ecogenics GmbH (Balgach, Switzerland). Size-selected fragments of genomic DNA were enriched for SSR content with magnetic streptavidin beads and biotin-labelled GATA, GTAT, AAAC and AAAG repeat oligonucleotides. SSR-enriched library was sequenced on an Illumina MiSeq platform using the Nano 2x250 v2 format. After assembly, 3,855 contigs or singlets contained a microsatellite insert with a tetra- or a trinucleotide of at least six repeats or a dinucleotide of at least 10 repeats. Primer design was possible in 171 microsatellite candidates. Out of these, 11 loci were successfully amplified in all 15 screened individuals. For additional testing, we further amplified these 11 loci in 32 individuals of *L. cymosa* on an automatic capillary sequencer (MegaBACE 1000, GE Healthcare, Chicago, IL, USA) with the size standard MegaBACE ET400-R (GE Healthcare, Chicago, IL, USA).

We assembled the primers for the targeted microsatellites in multiplexes according to their annealing temperatures (Table 2).

Table 2 Description of genetic markers for each microsatellite loci, annealing temperature (Ta) and Genbank accession numbers.

| Primer | Primer sequences (5’- 3’) | Repeat motif | Size (bp) | Ta (°C) | GenBank accession no. |
| --- | --- | --- | --- | --- | --- |
| Leo80 | F: TTTAGCGGTACGCTTTTCAC  R: AAAAGCATGGCCTTTCCAGC | (TTGT)7 | 224-236 | 53 | MF002374 |
| Leo89 | F: GTTCGCCTCACCATAAAGGC  R: AAGAGTGAGCATGCGTGAAG | (TTTC)8 | 197-221 | 55 | MF002375 |
| Leo94 | F: AAACCCTTGTTTTCGAATTTAGATG  R: GGGGCCAATTTGACTTTTTGC | (TTTG)8 | 220-236 | 59 | MF002376 |
| Leo270 | F: GTACTTGCACCATGCCACC  R: TAGCACTTCTGCACTTGTTG | (AAAC)8 | 110-122 | 55 | MF002377 |
| Leo466 | F: AGCATAGACACCACGGCTAC  R: AACTTGATCCCCAGTTTGGC | (AAGA)9 | 196-216 | 55 | MF002378 |
| Leo1842 | F: ACCCCATGACCCTTTAGTGC  R: TTTTATGTTAAGTTCTTGCAATGGG | (AAAG)7 | 224-244 | 59 | MF002379 |
| Leo2254 | F: ATGCACCATTGAACTTGGTC  R: AACCCACGCCTTTTATGCAG | (AAGA)8 | 126-166 | 53 | MF002380 |
| Leo2428 | F: TTATATTTGTCCTCCCTTCTGATAAC  R: GATCAATGGCTGCTCTCGTG | (TTTG)7 | 100-112 | 59 | MF002381 |
| Leo2433 | F: AGGAGTTAGCAATACAAAGTGAGTG  R: TCGTGTTAATCCCTTCTTTCCC | (ATAC)14 | 216-268 | 59 | MF002382 |
| Leo2833 | F: ACTATGTCACCTCACAAGCC  R: CTGAAATGCACCCTACGGAAC | (CATA)8 | 178-206 | 53 | MF002383 |
| Leo2853 | F: TTGCAAGGCACAATGACGAC  R: TACACAGTGCCAACATGCAG | (ATAC)8 | 158-190 | 55 | MF002384 |

We performed PCR reactions using the Qiagen Type-it microsatellite PCR kit (Qiagen, Venlo, Netherlands) in 14.6 µl of master solution containing: 2 µl of 10 ng/µl genomic DNA, 8.3 µl 1x Type-it multiplex PCR master mix, 1.2 to 2.0 µl of each 2 µM primer pair solution (based on calibration), and double-distilled water (ddH_2_O). PCR conditions on the thermocycler (T1, Biometra, Goettingen, Germany) were set to 94 °C for 5 min, followed by 34 cycles at 94 °C for 30 s, annealing temperature for 90 s, extension at 72 °C for 30 s and a final extension at 60 °C for 30 min (Table 2). PCR amplification products were again separated on the MegaBace 1000. Subsequently, we determined the allelic composition of each sample using the MegaBACE Genetic Profiler v. 2, an example of an electropherogram for each primer is given in Figure 1.

Figure 1 Example of individual electropherograms for multiplexes 1-3 (a-c). Peaks are colored according to the dyes used for each primer (FAM: blue, HEX: green, TMR: black) and marker ranges are indicated by vertical bars on top of the respective peaks

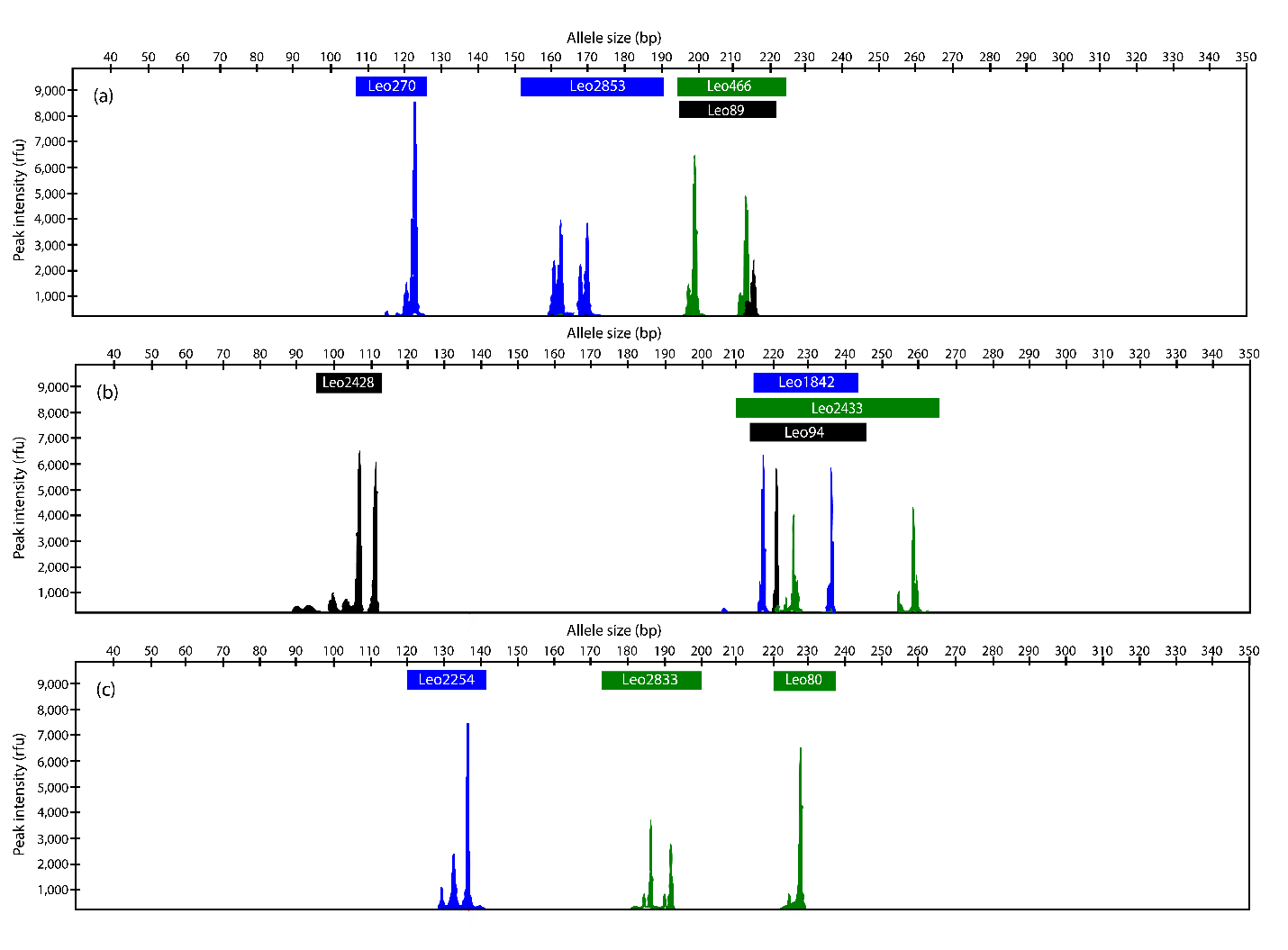
***Population genetics analysis***

Effective number of alleles (A_E_), observed heterozygosity (H_O_), expected heterozygosity (H_E_), and deviation from Hardy-Weinberg equilibrium (HWE) were determined using GenAlex v. 6.2 (Peakall and Smouse, 2006) based on 645 genotyped *L. cymosa* individuals from EBQB. Linkage disequilibrium (LD) was tested using GENEPOP version 4.3 (Rousset, 2008). Null alleles were analysed using MICRO-CHECKER v. 2.2.3 (Van Oosterhout et al., 2004).

**Results**

Amplification products of 11 microsatellite primer pairs showed polymorphic bands that could be reliably scored and were used for further analysis. Mean number of alleles of all loci was 5.7 (range 3 – 14) and expected heterozygosity ranged from 0.12 to 0.87 (mean 0.41). Significant deviations from HWE were found in five loci (Table 3). Significant linkage disequilibrium (q-value < 7 x 10^-5^) was detected for Leo89 and Leo466 (Storey and Tibshirani, 2003). Null alleles were detected for Leo2428 (A_N_= 0.053). However, corrected estimated allele frequencies based on null alleles deviated only by 0.02 from the observed frequencies (Table 3).

Table 3 Number of alleles (A_N_), Effective number of allele (A_E_), Observed heterozygosity () and expected heterozygosity (), HWE deviations (P < 0.05 = *, P < 0.01 = **, P < 0.001 = ***)

| *L. cymosa* [EBQB] (n = 664) | | | | |
| --- | --- | --- | --- | --- |
| Locus | **A** | **A_E_** | **H_o_** | **H_e_** |
| Leo80 | 3 | 1.9 | 0.423 | 0.469** |
| Leo89 | 6 | 1.4 | 0.307 | 0.307 |
| Leo94 | 3 | 1.1 | 0.117 | 0.119 |
| Leo270 | 4 | 1.1 | 0.116 | 0.125** |
| Leo466 | 4 | 1.9 | 0.488 | 0.484 |
| Leo1842 | 4 | 2.1 | 0.495 | 0.514* |
| Leo2254 | 8 | 2.7 | 0.660 | 0.632*** |
| Leo2428 | 4 | 1.5 | 0.317 | 0.349 |
| Leo2433 | 14 | 7.3 | 0.817 | 0.863*** |
| Leo2833 | 6 | 2.6 | 0.562 | 0.609 |
| Leo2853 | 7 | 1.4 | 0.256 | 0.271 |

All microsatellite markers were successfully amplified in the second population of *L. cymosa* and congeneric species *L. glycicarpa.* While seven loci out of 11 were successfully amplified in *L. crassa* (Table 4). The microsatellite markers where highly variable in the population of *L. cymosa* at EBQB and cross-amplification maintained this high variability. For all sampling sites, fragment length was within the same range as in *L. cymosa* from EBQB.

Table 4 Amplification success among *Leonia cymosa* populations and congenerics

| Loci | L. cymosa [EBQB]  (n = 664) | L. cymosa [AM]  (n = 6) | L. crassa [AM]  (n = 3) | L. glycicarpa [AM]  (n = 5) |
| --- | --- | --- | --- | --- |
| Leo80 | + | + | + | - |
| Leo89 | + | + | + | + |
| Leo94 | + | + | + | - |
| Leo270 | + | + | + | - |
| Leo466 | + | + | + | + |
| Leo1842 | + | + | + | - |
| Leo2254 | + | + | + | + |
| Leo2428 | + | + | + | + |
| Leo2433 | + | + | + | + |
| Leo2833 | + | + | + | + |
| Leo2853 | + | + | + | + |

**Supporting Information – S2**

**R script for executing function for extracting linear travel distances from movement data and for executing the CMG method (combination of movement data and gut passage time).**

####Linear.distances() function

linear.distances <- function (time, year, month, day, xUTM, yUTM){

timeN <- sapply(strsplit(time,":"),

function(x) {

x <- as.numeric(x)

x[1]+x[2]/60#+x[3]/1200

} ) #converts time to decimal (e.g 11:30=11.5, 01:00 = 1.0)

time_interval <- abs(apply(combn(timeN,2), 2, diff))

year_interval <- abs(apply(combn(year,2), 2, diff))

month_interval <- abs(apply(combn(month,2), 2, diff))

day_interval <- abs(apply(combn(day,2), 2, diff))

X_interval <- abs(apply(combn(xUTM,2), 2, diff))

Y_interval <- abs(apply(combn(yUTM,2), 2, diff))

distance_interval <- sqrt((X_interval^2)+(Y_interval^2))

date_interval <- year_interval+month_interval+day_interval

comb <- data.frame (date_interval, distance_interval,time_interval)

daily_linear_travel_paths <- comb[ which(comb$date_interval == 0), ]

daily_linear_travel_paths$date_interval <- NULL

return(daily_linear_travel_paths)

}

#example of input parameteres

time <- as.character(c("11:00", "11:30", "12:00", "12:30", "13:00", "13:30"))

year <- as.numeric(c(2012,2012,2012,2012, 2012))

month <- as.numeric(c(12,12,12,12, 12, 12))

day <- as.numeric(c(14,14,14,14,14,14))

xUTM <- as.numeric(c(704265,704256, 704249, 704146, 704090, 704010))

yUTM <- as.numeric(c(9517640, 9517554, 9517526, 9517567, 9517564, 9517571))

##########to obtain SDD estimates using the CMG method#######

##1. Load data file from csv, time format should be in "%H:%M” or "%H:%M:%S", and date should be separated in columns according to day, month, year.

trial <- read_csv("~/linearmovement_automatization_trial.csv",

locale = locale(date_format = "%Y-%m-%d",

time_format = "%H:%M:%S",tz = "UTC"))

##2. Restrict data to fruiting season

trial[trial$Month %in% c("3","4","5"),]-> trial_FS

##3. Order data chronologically

trial_FS <- trial_FS [order(trial_FS$Year, trial_FS$Month, trial_FS$Day, trial_FS$Time),]

\## 4. Determine input parameters for function

as.character(trial_FS$Time) ->time #format "%H:%M" if "%H:%M:%S" then add +x[3]/1200 to function by deleting “#” on line 5.

as.numeric (trial_FS $Year )-> year

as.numeric(trial_FS$Month) -> month

as.numeric(trial_FS$Day) -> day #data points have to be in chronological order

as.numeric(trial_FS$X) -> xUTM

as.numeric(trial_FS$Y) -> yUTM

##5. execute function

linear.distances (time, year, month, day, xUTM, yUTM) -> daily_linear_travel

##6. restrict linear travel paths to those within the gut passage time of a particular plant species or a mean for the animal species.

CMG_SDDestimates <-daily_linear_travel[daily_linear_travel$time %in% c(1,1.5,2),] #e.g. gut passage of 1-2hrs

**Supporting Information – S3**

**Supporting Figures**

**Supporting Figure S3.1:** Examples of daily travel paths of a mixed species group of tamarin monkeys, *Saguinus mystax* and *Leontocebus nigrifrons*. Each line represents a daily travel path recorded in 2013, with some examples highlighted with different colours. On average, tamarins cover 1508.9 ± 251.8 m per day.

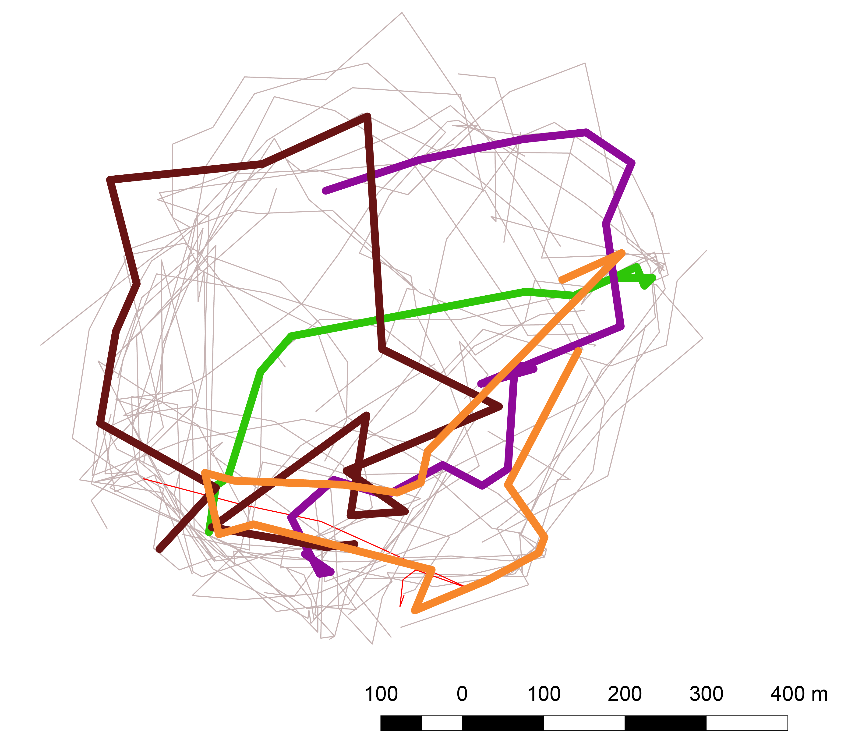

**Supporting Figure S3.2:** Map of sampled individuals of *Leonia cymosa* (seedlings ◊, juveniles ○ and adult trees ⌂) within quadrats, collected in 2014 (dark grey) and collected in 2015 (light grey). Transects for sampling are depicted with dashed lines. Home ranges of the tamarin groups 1 and 2 are marked with solid grey lines.

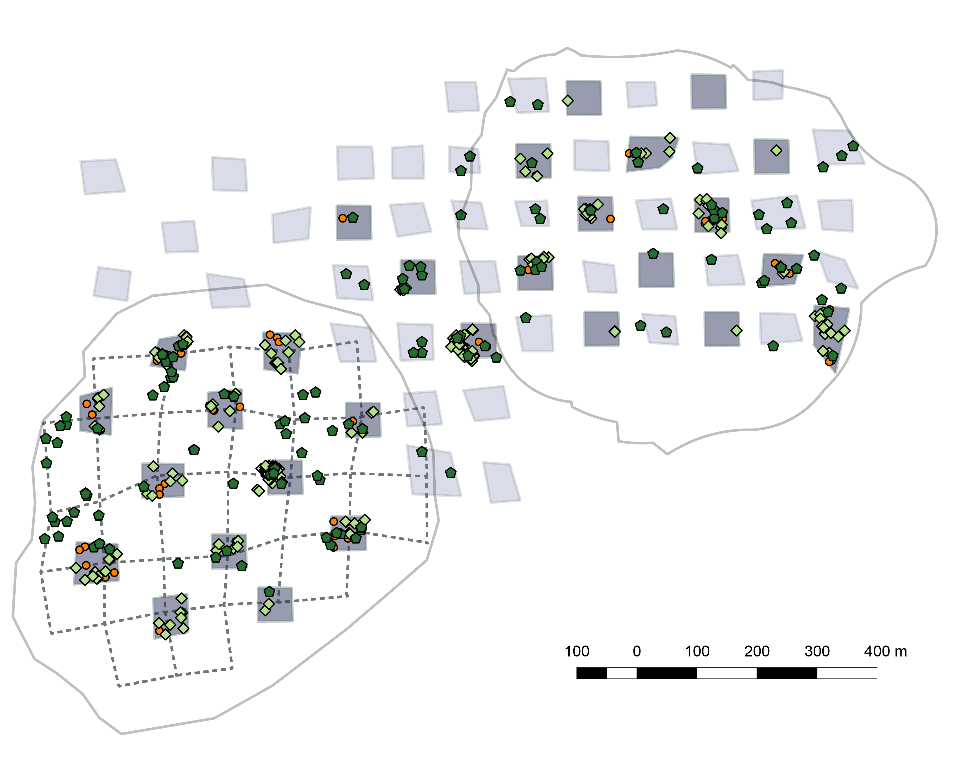

Group 2

Group 1

**Supporting Figure S3.3:** Sampling map for *Parkia panurensis* (seedlings ◊, juveniles ○ and adult trees ⌂) Walking pathways are depicted with dashed lines. Home ranges of the tamarin groups 1 and 2 are marked with solid grey lines.

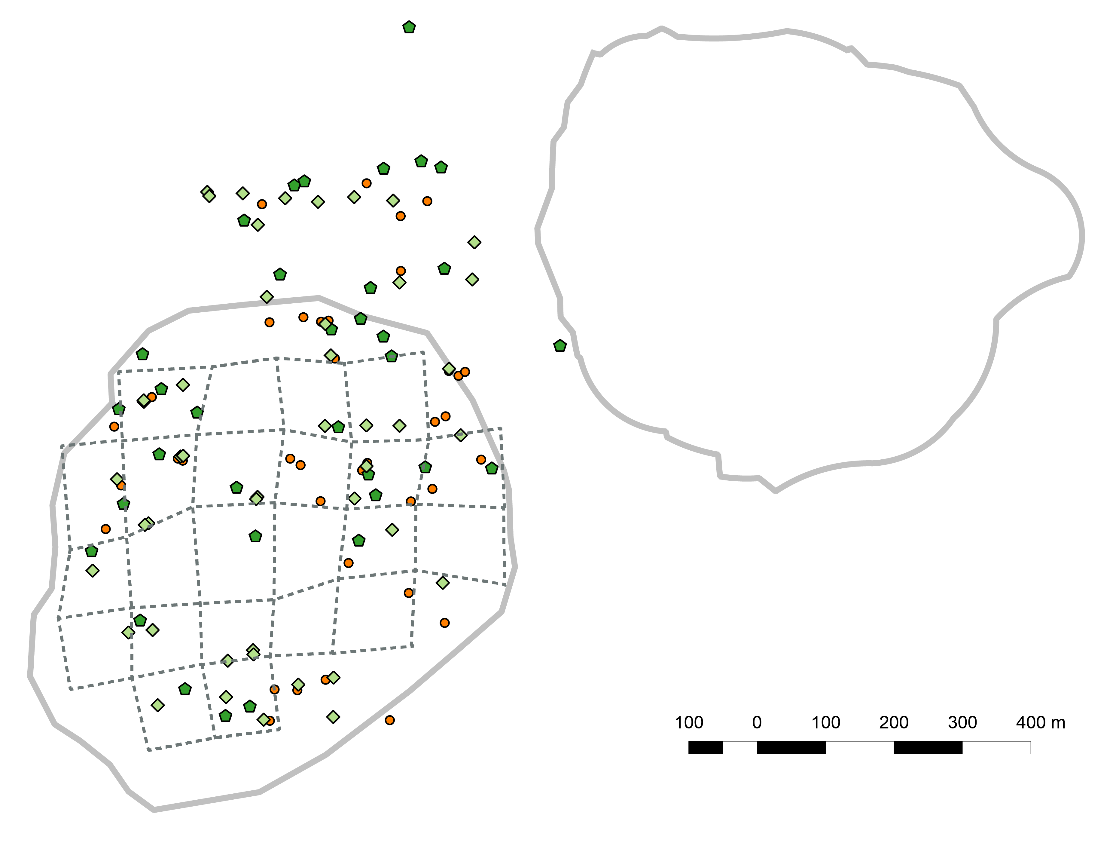

Group 1

Group 2

Figure S.3.4. Cumulative distribution of SDD for *Parkia panurensis* (A) and *Leonia cymosa* (B) for the five approaches used in this study: Observed seed dispersal events (OSD), genotyped seed coats (GSC), parental analysis fo seedlings (PAS), combination of movement data and gut passage (CMG), and individual-based modelling (IBM)

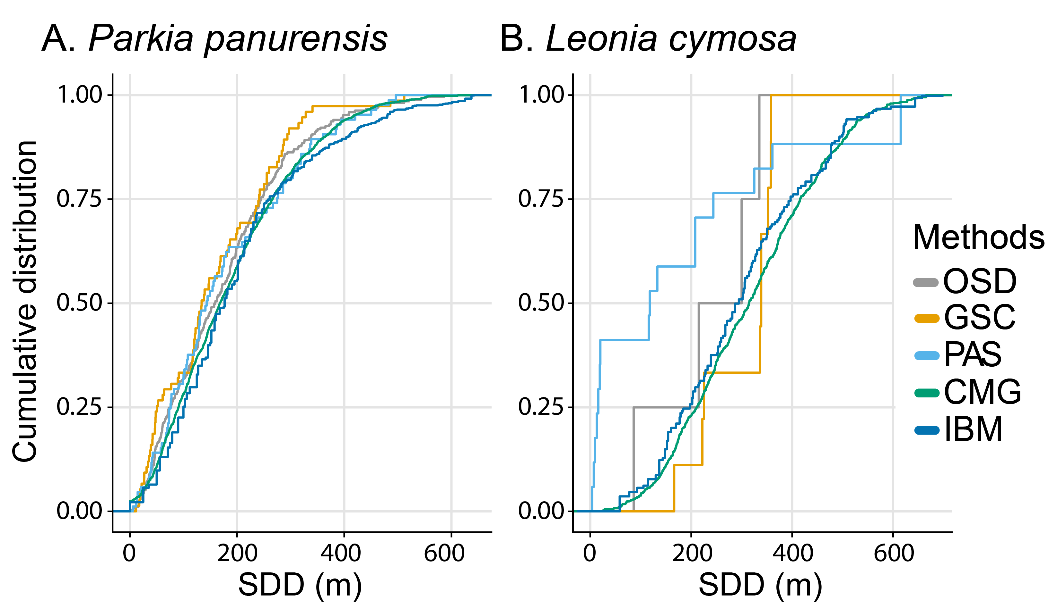

Figure S.3.5. Comparison of SDD estimates between species for each method used in this study: observed seed dispersal events (OSD), genotyped seed coats (GSC), parental analysis of seedlings (PAS), combination of movement data and gut passage (CMG), and individual-based modelling (IBM) Horizontal lines represent medians, boxes the 25-75% quartiles, dots are outliers. Bars above the boxplots indicate differences between species for each method based on Wilcoxon rank sum test.

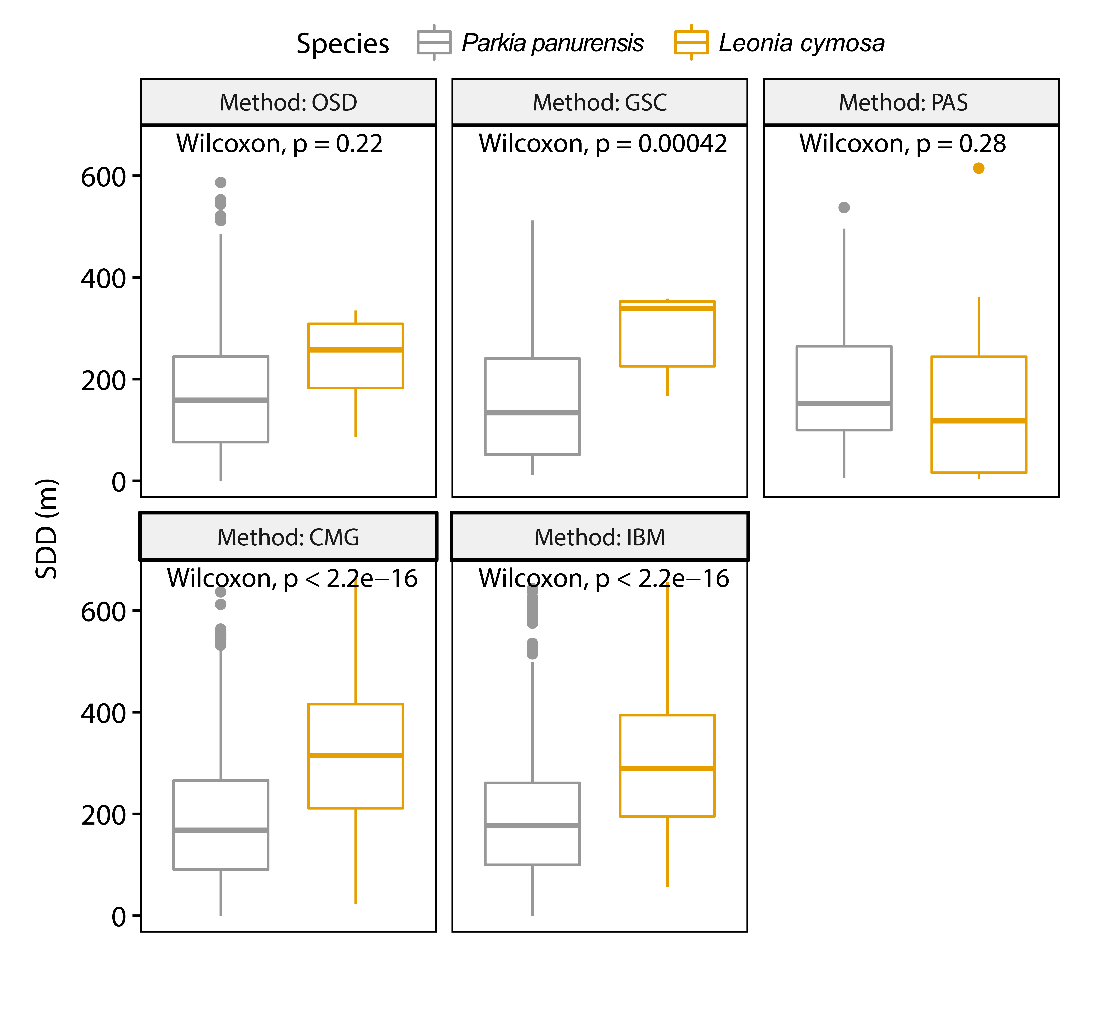

**Supporting Information – S4**

**Supporting Tables**

Table S4.5: Parameter and values used in the simulation of the individual-based model method (IBM)

| Name | Value | Comment |
| --- | --- | --- |
| no_cells | 80 | size of the landscape each direction |
| scaling | 25 | size of on cell |
| sim_time | 108 | def. time 580 min = 9:40 +/-x hours/day |
| no_days | 100 | no of simulated days = 100 days |
| start_tree_x | 704 100.34 | 704224.39 |
| start_tree_y | 9 517 409.65 | 9517563.63 |
| tree_file | all_trees_with_leonia-41.txt | with 41 Leonia trees |
| start_energy | 60 | starting energy at day 1 every simulation |
| energy_level_1 | 80 | if below, feeding |
| energy_level_2 | 150 | if above, other behaviour |
| tree_time | 0.1 | change tree – ̃ 5 time steps |
| tree_time_leonia | 0.25 | change tree – ̃ 2 time steps |
| feeding | 8 | gain during feeding per time step |
| feeding_leonia | 4 | gain during feeding per time step |
| running | −1.6 | loss during running per time step |
| marking | −1.7 | loss during marking per time step |
| foraging | −0.5 | loss during foraging per time step |
| resting | −0.7 | loss during resting per time step |

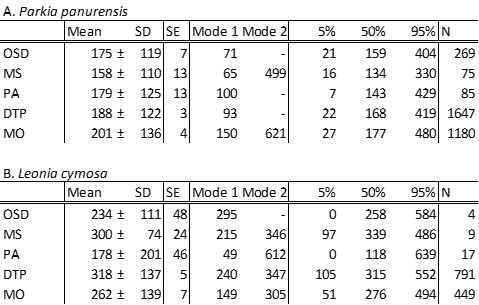

Table S4.2 Summary of seed dispersal distance estimates for *Leonia cymosa* (A) and for *Parkia panurensis* (B) using different methods (OSD= observed seed dispersal events, MS = Maternal ID from seed coats, PA = Parentage analysis, DTP = from daily travel paths, MO = from individual-based modelling)
